# CRISPR/Cas-based genome editing for cyanophage of *Anabeana sp*

**DOI:** 10.1101/2024.04.16.589687

**Authors:** Shengjian Yuan, Yanchen Li, Chunhua Kou, YiChen Sun, Yingfei Ma

## Abstract

Efforts have been directed towards genome editing in cyanobacteria, yet achieving genome reduction in cyanophages remains a challenging task. In this study, we utilized the CRISPR-Cas12a system to successfully delete multiple genes within A-1(L) and A-4(L) cyanophages. Through careful manipulation, we generated a deletion mutant cyanophage with a 2,778 bp reduction in genome size, representing a 6.6% decrease compared to the wild type (WT). In summary, our research has introduced a robust method for gene editing in cyanophages, facilitating the identification of nonessential genes essential for cyanophage propagation. This advancement holds promise in addressing the widespread issue of water blooms and the associated environmental hazards.

## Introduction

Water blooms are a global issue that occurs seasonally and is primarily caused by the eutrophication of water bodies^1^, leading to the overgrowth of cyanobacteria such as *Aneabena, Microcystis, Aphanizomenon, Planktothrix*, and others^2^. These blooms have detrimental effects on freshwater quality and ecosystem health^3^. The release of toxins and metabolites by cyanobacteria during harmful algal blooms (CyanoHABs) poses a threat to the safety of human and animal life^4-6^. These CyanoHABs can produce hepatotoxic, neurotoxic, or dermatotoxic cyanotoxins^7^, with microcystins being a prominent hepatic toxin^8^. Other cyanotoxins like anatoxin-a or saxitoxins have strong neurotoxic effects, posing a risk to both animals and humans^9-11^. Current methods for treating water blooms primarily involve physical techniques like sonication and salvage^12^, the addition of chemical components such as algaecides (e.g., copper sulfate, hydrogen peroxide)^13^, or the use of microorganisms^14^. However, these methods have limitations in their effectiveness^15^. Therefore, there is an urgent need to develop new and efficient approaches to address water blooms.

Cyanophages, a type of phage that specifically infects cyanobacteria, have shown potential as an effective tool for treating water blooms^16^. However, the wild-type cyanophages have limitations in their application due to their host-specificity and resistance^17^. Currently, research on cyanophages primarily focuses on their isolation, characterization, and genome sequencing^18^, with limited studies on gene editing. Phage engineering has made significant advancements^19^, including enhancing antimicrobial activity, expanding host range^20^, incorporating fluorescence visualization, and fusing functional genes. These engineering techniques can also improve the application of cyanophages, expanding our understanding of cyanophages biology and their potential uses in combating water blooms.

The CRISPR system has already been utilized in phage genome engineering^21, 22^. However, due to the significant differences between cyanobacteria and bacteria, there is limited information available regarding genetic tools for editing cyanophage genomes. Considering previous studies demonstrating the toxicity of Cas9 protein to *Anabeana* sp. PCC 7120, the Cas12a system was chosen for editing cyanobacterial genomes^23^. This system has successfully enabled the insertion of green fluorescence protein into the *Anabeana* 7120 genome and the knockout of scarfford genes within the same genome^24^. Based on these findings, our objective is to manipulate cyanophage genomes using the Cas12a system.

The CRISPR-Cas12a system, which belongs to class 2 systems, type V, is a type of CRISPR protein. It only requires a single guide RNA to recruit the Cas12a protein and bind to the target sequence^25^. The target sequence is typically located 19-23 nucleotides downstream of the PAM sequence (TTN)^26^. Upon binding, the Cas12a protein induces staggered cuts, resulting in double-strand breaks (DSBs)^27, 28^. Therefore, we aim to construct a platform for editing cyanophage genomes. By utilizing gene knockouts, we can elucidate unknown gene functions and determine their essentiality. Additionally, genome reduction allows for the insertion of foreign genes, such as mlrA, which can reduce the presence of microcystins in the environment during cyanophage infection. Ultimately, this engineering system can be developed to uncover fundamental aspects of cyanophage biology.

In this study, we present a platform for identifying hypothetical or essential genes in cyanophages and for knocking out non-essential genes using a CRISPR–Cas12a system. Specifically, we applied this genetic engineering method to cyanophage A-1(L) and A-4(L) of Anabeana sp. PCC.7120, which belong to myoviridae and podoviridae family, respectively^27, 28^. This approach enabled us to determine the essentiality of genes in cyanophages. Importantly, we were able to use the gRNA library for large-scale gene manipulation. Our work establishes a Cas12a-based strategy for engineering cyanophages that could be broadly applicable. The use of cyanophages as anticyanobacterial agents requires a thorough understanding of phage biology and may offer a viable tool for addressing water blooms.

## Results

### Development of the Cyanophage editing approach

We utilized the CRISPR/Cas12a system derived from *Francisella novicida*^29^ for editing the cyanophage genome. The original pCpf1b-sp^24^ (180423N1) plasmid comprises Cpf1, RSF1010 replication origin, a spectinomycin-resistant gene, SacB gene, lacZ gene, and a gRNA array with three direct repeat (DR) sequences. In order to streamline the plasmid, redundant genes including the SacB gene, lacZ gene, and two direct repeat sequences were excised, resulting in the creation of pCpf1b-sp2 (Fig. S1).

Initially, we annotated the gene functions of cyanophage A-4(L) and A-1(L) by screening each ORF of its genome using BlastP in the NCBI non-redundant (Nr) database (Fig. S2 and Fig. S3). Based on this analysis, we postulated that certain predicted hypothetical genes within the A-4(L) cyanophage genome could be deleted. Next, specific guide RNAs (gRNAs) with a protospacer adjacent motif (PAM) sequence of TTN were selected and constructed the plasmid as outlined in the Methods section. The constructed plasmids were introduced into the *Anabeana* PCC 7120 strain using a conjugation system^28^ (Fig. 1A).

**Fig. 1.**
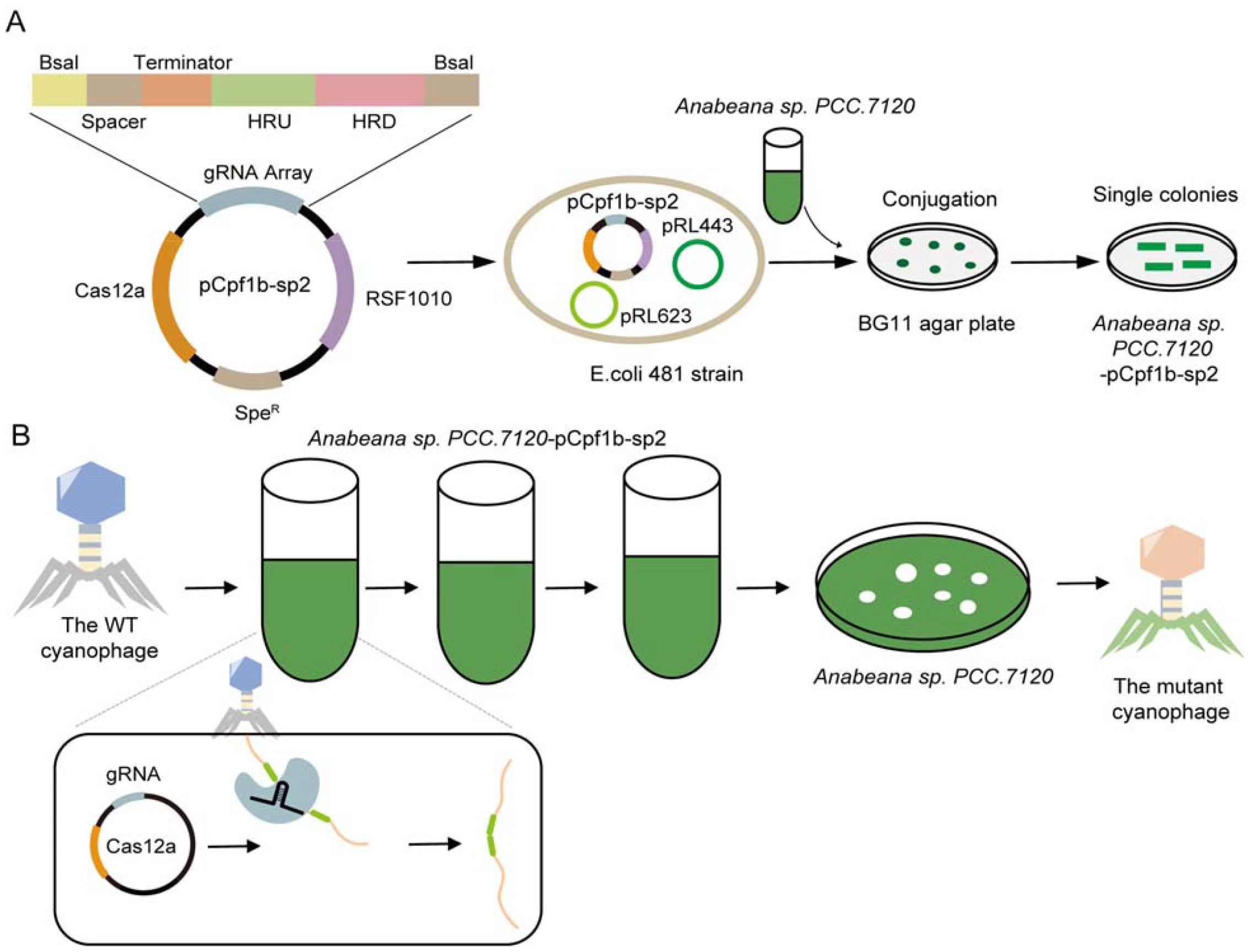
Workflow of CRISPR/Cas genome editing in cyanophage. **(A)** Detailed schematic illustrating the construction of CRISPR-Cas12a system. The yellow diamond represents the two BsaI sites used for ligating the backbone. The brown section represents the spacer sequences, selected from downstream of the PAM sequences. The orange section indicates the terminator of the gRNA array. Green and pink diamonds represent the 50 bp upstream and downstream sequences of the target gene, respectively. The plasmid was assembled using Golden Gate assembly and then transferred into *Anabeana* cells via conjugation. The exconjugants were streaked on BG11 agar plates supplemented with antibiotics. **(B)** Gene editing of cyanophages occurred during the transfer process. Mutant cyanophages were isolated using double-layer BG11 agar plates.

After confirming the presence of the target plasmid in the *cyanobacteria*, they were infected by a specific cyanophage. During cyanophage infection, the injection of cyanophage DNA into the cell triggers a process where the single-guide RNA (sgRNA) encoded by pCpf1b-sp2 activates the Cas12a nuclease (encoded by pCpf1b-sp2), leading to the cleavage of the gene at the desired site. Subsequently, the resulting break is repaired through homologous recombination (HR) induced by the HR sequence (pCpf1b-sp2), resulting in gene deletion^30^. If the deleted gene is non-essential for cyanophage propagation, the mutant cyanophage progeny can proliferate. This mutant cyanophage population is continually transferred to fresh cells containing the plasmid system, facilitating the accumulation of gene deletions in the cyanophage genome (Fig. 1B).

Cyanophage A-4(L) is a T7-like podovirus that infects *Anabeana sp. PCC 7120*. Its genome consists of 38 ORFs, with 14 predicted functional genes and the remainder being hypothetical genes. Additionally, the genome contains 810 bp repeats on both ends^27^. To assess the feasibility of this method, we specifically targeted *gp2* of A-4(L) for its short length of approximately 200 bp and its potential for knockout (Fig. 2A). Over multiple generations, the mutant cyanophage was isolated, and the deletion of *gp2* was monitored using PCR amplification. The results indicated that a 165 bp sequence of *gp2* was successfully knocked out (Fig. 2B). Sanger sequencing further confirmed the accuracy of the deletion. Subsequently, following the same protocol, we acquired another mutant cyanophage with a 315 bp knockout of *gp8* of A-4(L) (Fig. 2B). For *gp18* of A-4(L), initially, four gRNAs were selected with 25 bp downstream of the PAM (TTG) using the CHOPCHOP software^31^. One of these gRNAs produced a faint knockout band during cyanophage generation, but failed to isolate the mutant cyanophage due to its weak efficiency. Subsequently, we replaced it with three new gRNAs (sequences obtained from Juyuan Zhang, Institute of Hydrobiology, Chinese Academy of Science) for knocking out *gp18*. Furthermore, we optimized the gRNA by predicting secondary structure, as an optimal structure could enhance knockout efficiency (http://rna.tbi.univie.ac.at/). Following the same protocol, we identified a potent candidate gRNA and successfully obtained the mutant cyanophage with a 400 bp knockout of *gp18* (Fig. 2B).

**Fig. 2.**
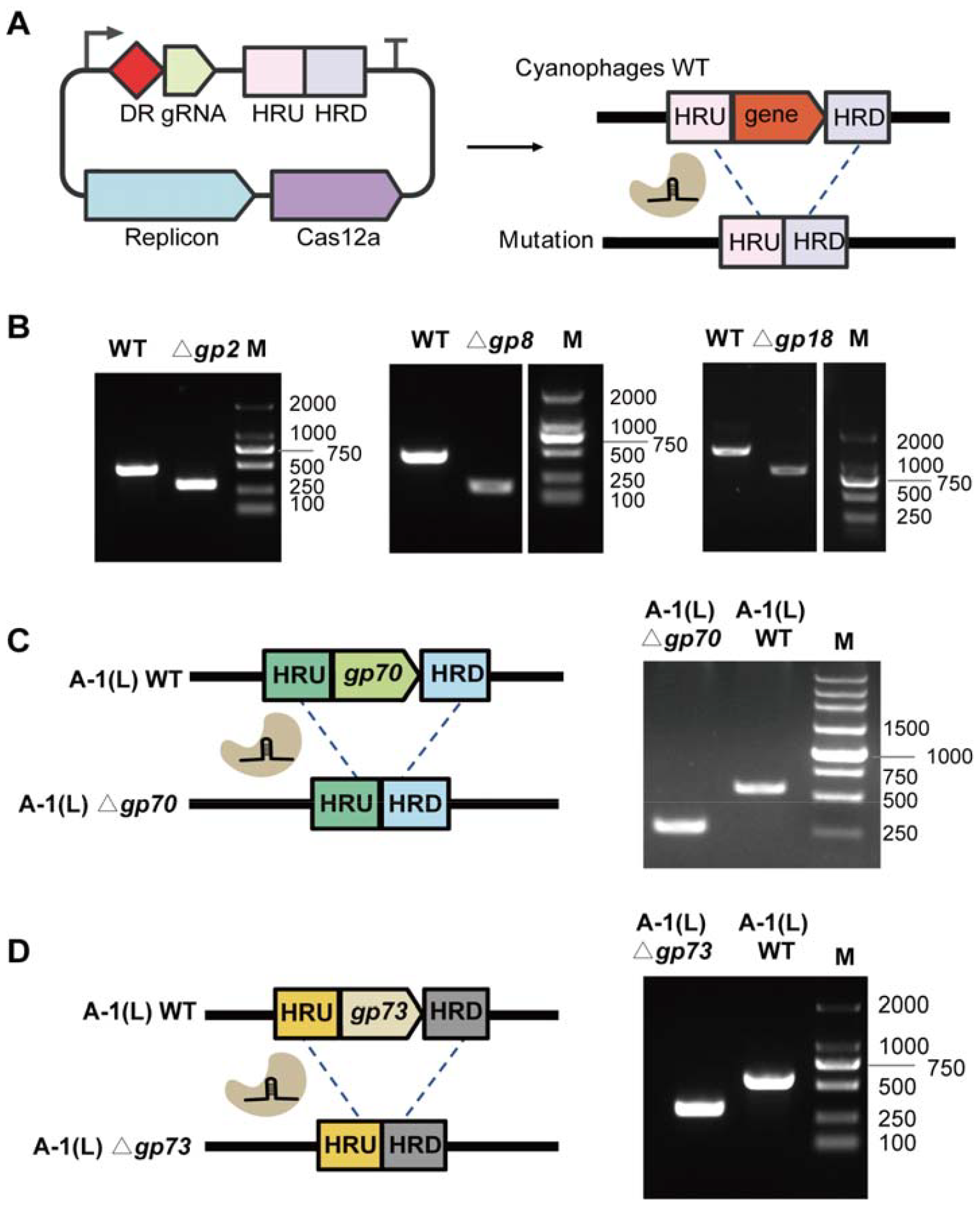
Single gene deletion using CRISPR-Cas12a system. **(A)** The plasmid was utilized for deleting genes in cyanophages. The red diamonds represent the spacer sequence, while the pink and light purple diamonds represent the homologous arms of the upstream and downstream regions, WT: wild-type, △g*p8*: Deleted *gp8*. **(B)** Gel electrophoresis results of A-4(L) gene deletions. All mutant cyanophages were confirmed through Sanger sequencing. **(C)** The design of *gp70* gene deletion in A-1(L) cyanophage and the gel electrophoresis results. **(D)** The design of the gp73 gene deletion in the A-1(L) cyanophage and the corresponding gel electrophoresis results of gene deletions.

Additionally, we manipulated the lytic myoviridae cyanophage A-1(L) using the CRISPR-Cas12a system. In our previous study, we conducted a BlastP analysis of the A-1(L) cyanophage genome using the NCBI Nr database and annotated the predicted hypothetical genes. Subsequently, we selected several of these genes for deletion experiments to assess their essentiality. Specifically, we successfully knocked out *gp70* and *gp73* (Fig. 2C, D). However, despite our efforts, we were unable to generate a mutant cyanophage for *gp66*. Next, we replaced *gp70* with GFP on the mutant cyanophage genome where *gp73* was deleted. Surprisingly, we observed the emergence of a smaller mutant cyanophage displaying a partial deletion of *gp70* and *gp71*. Consequently, we have concluded that this genome editing system is valuable for the A-4(L) and A-1(L) cyanophage, and further applications can be explored to investigate additional hypothetical genes within the A-4(L) and A-1(L) genome, providing us with valuable insights into cyanophage biology.

### Iterative knock-out and simultaneous editing through CRISPR-Cas12a system for cyanophage targeting

We successfully obtained an effective A4 *gp2* gRNA and demonstrated that *gp2* could be deleted. To assess the efficacy of other gRNAs and explore the possibility of simultaneous editing of two genes, we designed a plasmid with homologous arms selected from the upstream of *gp2* and the downstream of *gp3* (Fig. 3A). Following the generation, we isolated mutant cyanophages with a successful deletion of 871 bp encompassing *gp2-gp3* (Fig. 3B). To explore the potential for iterative knock-out in the cyanophage system, we aimed to delete *gp5* in the A-4(L)-?*gp2* knock-out mutant cyanophage. Two gRNAs were selected, and one of them was proven to be effective in deleting *gp5* (Fig. 3B). PCR and Sanger sequencing confirmed the successful generation of a mutant cyanophage with deletions in both *gp5* and *gp2*, totaling 389 bp (182+207 bp).

**Fig. 3.**
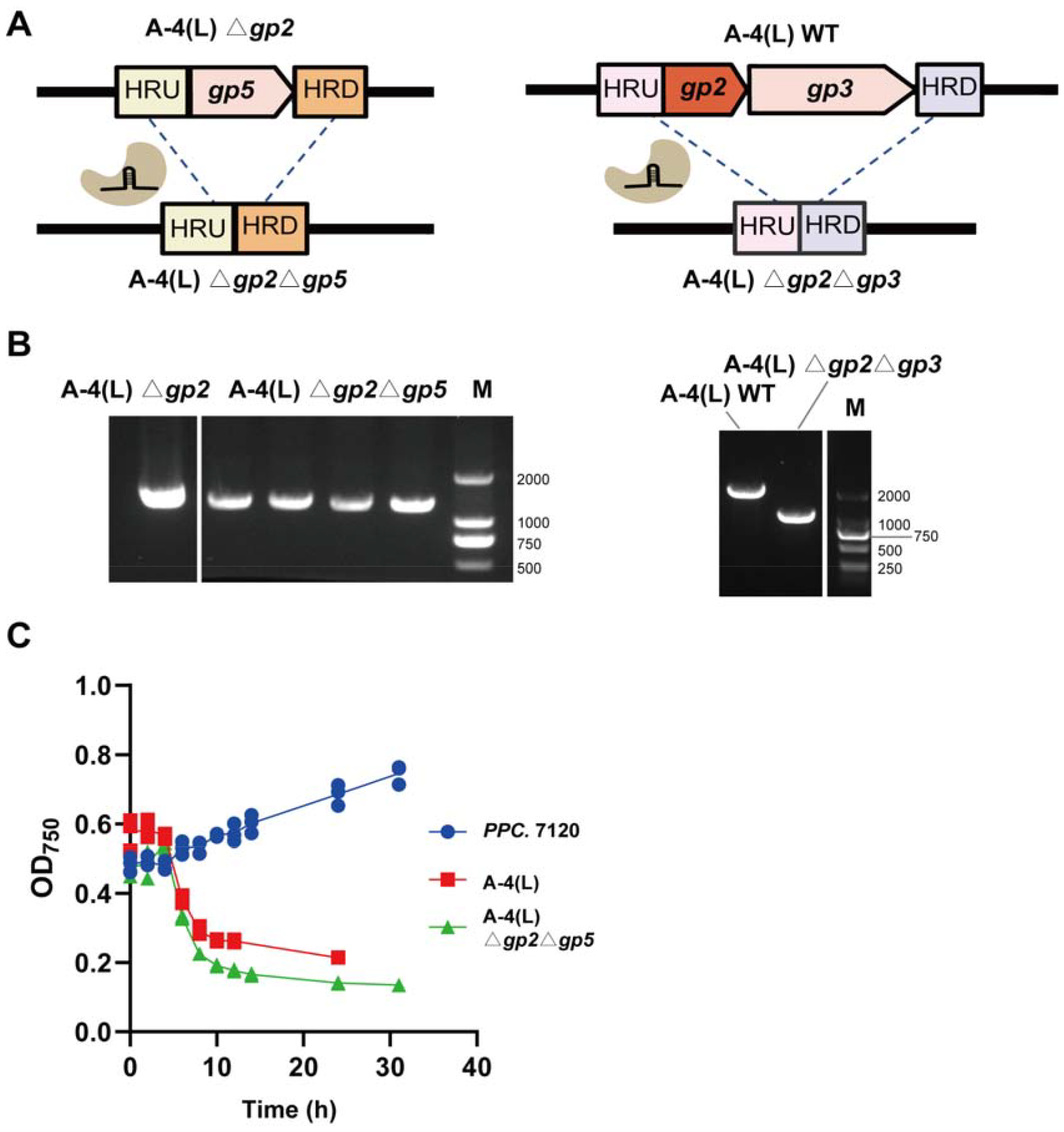
Multigene knock-out through the CRISPR-Cas12a System. **(A)** Overview of the multigene knock-out of A-4(L) cyanophage. **(B)** Gel electrophoresis results of gene deletions. M: 2000 bp marker. **(C)** Killing curves for *cyanobacteria* after infection with the A-4(L) mutants. The blue line represents *Anabeana* cell 7120 without infection of cyanophage. The red line represents the wild-type A-4(L), and the green line represents the A-4(L)-△*gp2*△*gp5* mutant cyanophage. **(D)** One-step growth curve of wild-type and mutant cyanophage. The green line represents WT A-4(L), the blue line represents A-4(L)-△*gp2*△*gp3*, and the pink line represents A-4(L)-△*gp2*△*gp5* mutant cyanophage. Data shown represent the mean ± standard deviation of the values obtained in three replicates.

The mutant A-4(L)-△*gp2*△*gp3* and A-4(L)-△*gp2*△*gp5* is viable phages that produced plaques of similar size and efficiency. Plaque assays indicated that the host ranges and plaque morphologies of both mutants closely resembled those of the WT phages. All variants demonstrated a strong ability to infect cyanobacteria and effectively inhibit their growth (Fig. 3C). Although there were slight differences in burst size among them, this variation is expected due to the differences in genome length.

So far, we have successfully deleted a variety of genes in the A-4(L) cyanophage, including *gp2, gp3, gp5*, and *gp18*. These gene deletions have resulted in mutants with varying burst sizes, yet all have displayed significant inhibitory effects on cyanobacterial growth. Importantly, our experiments have shown that the deletion of these genes does not disrupt the cyanophage life cycle.

### Generation and Characterization of the Smallest Genomes of A-4(L)

Next, we aim to knock out longer gene fragments in the A-4(L) genome to determine the extent to which genes can be deleted and seek to understand the genome capacity of A-4(L). Using the same gRNA that targets *gp2-gp3* of A-4(L), we have attempted to knock out *gp2-6, gp2-7*, and *gp2-8* by designing upstream and downstream homologous arm sequences at both ends of *gp2*-*gp6, gp2*-*gp7*, and *gp2-gp8*, respectively (Fig. 4A). These deletions would remove 1600 bp, 1826 bp, and 2078 bp, respectively, from the A-4(L) cyanophage genome, representing a maximum knockout fragment of nearly 5.6% of the complete genome (Fig. 4B). This marks the smallest mutant cyanophage of A-4(L) observed thus far and represents the first instance of cyanophage engineering using the CRISPR-Cas12a system.

**Fig. 4.**
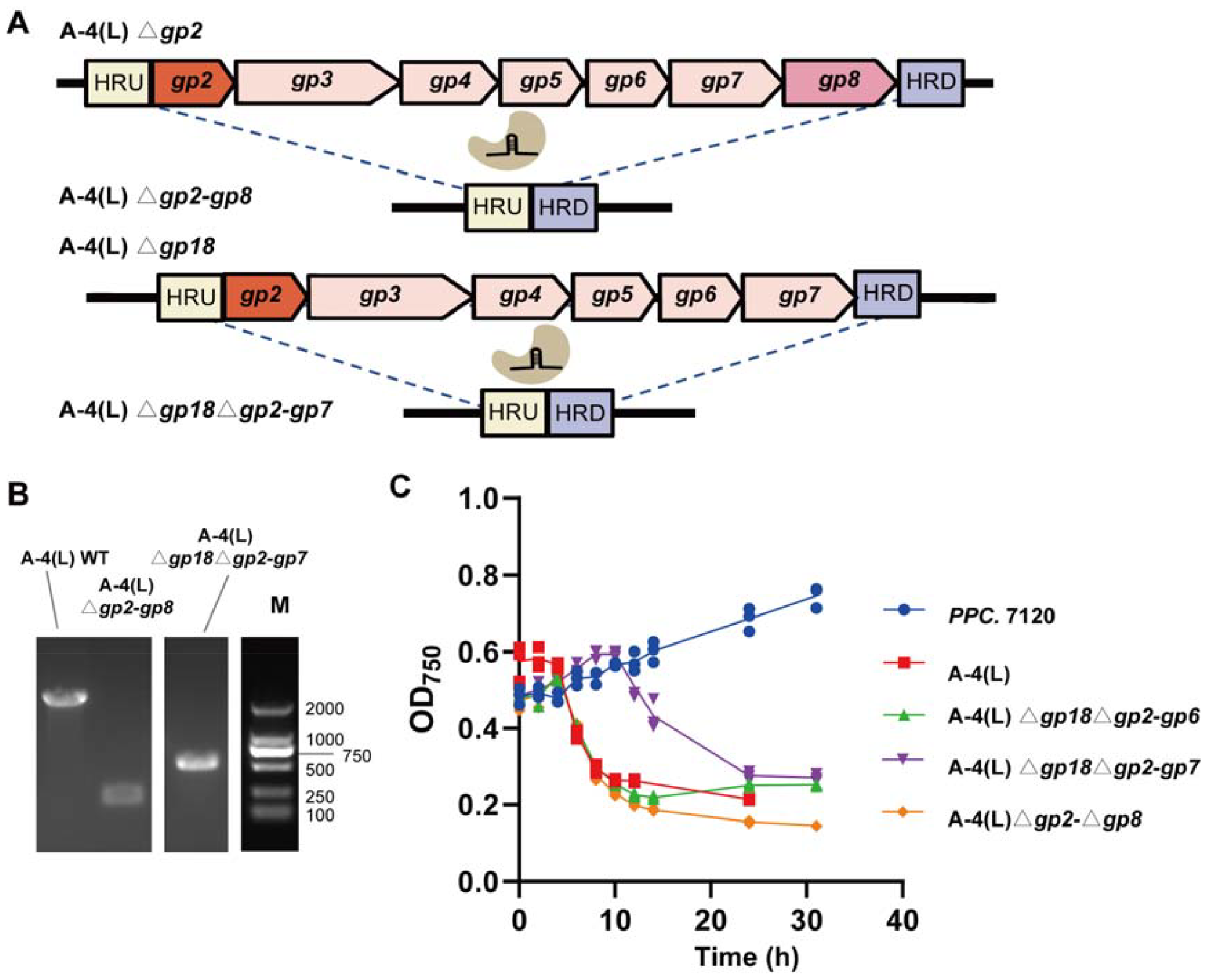
Deletion of large fragments by CRISPR-Cas12a system. **(A)** Overview of the large fragments deletion of A-4(L) cyanophage. **(B)** Gel electrophoresis results of gene deletions. M: 2000 bp marker. **(C)** Killing curves for *cyanobacteria* after infection with the A-4(L) mutants. The blue line represents *Anabeana* cell 7120 without infection of cyanophage. The red line represents the wild-type A-4(L), the orange line represents the mutant A-4(L)-△*gp2-*△*gp8* cyanophage, the purple line represents the A-4(L)-△*gp18*△*gp2-*△*gp7* mutant cyanophage, and the green line represents the A-4(L)-△*gp18*△*gp2-*△*gp6* mutant cyanophage. **(D)** One-step growth curve of wild-type and mutant cyanophage. The green line represents WT A-4(L), and the orange line represents the A-4(L)-△*gp2-*△*gp8* mutant cyanophage. Data shown represent the mean ± standard deviation of the values obtained in three replicates.

After isolating the A-4(L)-△*gp2-gp8* mutant phage among the three knock-out cyanophages, we were surprised to find that this mutant cyanophage could infect *Anabeana*. Furthermore, it was particularly surprising that the A-4(L)-△*gp18* knock-out mutant cyanophage continued to knock out *gp2-6* and *gp2-7* genes consecutively (Fig. 4B). This outcome demonstrates the effectiveness of our system for knocking out cyanophage genes and also allows us to identify hypothetical genes in the A-4(L) genome. The varying lengths of the deleted fragments suggest that genome capacity plays a crucial role in the survival of the cyanophage. This insight leads us to speculate that it could serve as a protective mechanism for the cyanophage progeny.

## Discussion

In this study, we aimed to investigate the effects of CRISPR/Cas12-mediated genome editing on two cyanophages, namely A-4(L) and A-1(L). Utilizing the CRISPR/Cas12 system, we successfully reduced the genome sizes of these cyanophages, resulting in the generation of a diverse population of genome-reduced cyanophages. Notably, we achieved a significant 6.2% reduction in the genome size of A-4(L).

Interestingly, among the diverse population of genome-reduced cyanophages, we observed that mutants with deletions encompassing the *gp18* and *gp2-gp5* genes, as well as those with deletions affecting the *gp2-gp8* genes in A-4(L), exhibited minimal impairment in infectious efficiency towards Anabeana. However, it is noteworthy that the simultaneous deletion of *gp18* and *gp7* in A-4(L) seemed to adversely affect its fitness. These genes are classified as hypothetical proteins and could potentially share similar functions. The loss of these two genes significantly impacts cyanophage propagation.

These findings highlight the potential impact of genome editing on cyanophage biology and shed light on the role of specific genes in cyanophage-host interactions. By manipulating the cyanophage genome, we can gain insights into the factors influencing their infectivity and replication dynamics. Further exploration of these observations could lead to practical applications in the field of environmental microbiology and contribute to the development of strategies for combating harmful algal blooms.

## MATERIALS AND METHODS

### Cyanobacteria, cyanophages, and media

The *Anabeana* sp. PCC.7120 used in this study was obtained from the laboratory of Chengcai Zhang at the Institute of Hydrobiology, Chinese Academy of Sciences. The host strain was cultured in BG11 broth^32^ supplemented with 30 °C temperature and a light intensity of 30 μE/m^2^/s. Alternatively, for solid media, 1.2% BG11 agar was used. The cyanophages A-1(L) and A-4(L) were obtained from the laboratory of Xudong Xu, also at the Institute of Hydrobiology, Chinese Academy of Sciences.

### Plasmid construction

The plasmid pCpf1b-sp^24^ (180423N1) used in this study was provided by the laboratory of Chengcai Zhang at the Institute of Hydrobiology, Chinese Academy of Sciences. For modifying the pCpf1b-sp plasmid, two fragments (7482bp, 4482bp) were amplified using Phusion DNA polymerase (New England BioLabs, M0530) via Polymerase Chain Reaction (PCR). Following gel electrophoresis, each fragment was purified using a Gel Extraction Kit (Omega, D2500-02). These purified fragments were then assembled using the Gibson assembly reaction (New England BioLabs, E2611L). The resulting plasmid, named pCpf1b-sp2, was utilized for cyanophage gene editing.

For gene delete in cyanophage, two amplification fragments were generated from the pCpf1b-sp2 plasmid with an additional Bsa I site modification. The N20 sequence was identified with a 22 bp region downstream of the PAM (TTN). The gRNA cassette was designed to include the gRNA, a terminator sequence, two 50 bp homologous arms (upstream and downstream) of the targeted genes, and two Bsa I sites. The gRNA cassette was obtained through overlapping PCR using three long synthetic oligonucleotides.

The two fragments from pCpf1b-sp2 plasmid and the gRNA cassette obtained above was assembled by using Golden Gate Assembly (Golden Gate Assembly Kit, New England BioLabs, and E1601S). The cycling conditions were as follows: 20 cycles of 37 °C for 3 min, 16 °C for 2 min, 60 °C for 30 min, 80 °C for 10 min, followed by holding at 4 °C. The recombination products were then transformed into DH5α electrocompetent cells. Subsequently, the cells were plated on Luria-Bertani (LB) 1.2% agar with 50 μg/ml spectinomycin and incubated at 37 °C. Colonies were verified by PCR and Sanger sequencing. The plasmids were extracted from the colonies using the TIANprep Mini Plasmid Kit (Tiangen, DP103).

### Transfer of Plasmid into Cyanobacteria by Conjugation

Conjugation was widely used to introduce foreign DNA into *Aneabena* sp. PCC 712032, following a previously established procedure^33, 34^. The *E. coli* 481 strain used for conjugation in this study was obtained from Xudong Xu at the Institute of Hydrobiology, Chinese Academy of Science. This strain carries a helper plasmid (pRL443) and a conjugative plasmid (pRL623). For transformation, the editing plasmids were introduced into competent cells of the 481 strain grown in LB broth supplemented with 25 μg/mL ampicillin, 12.5 μg/mL chloramphenicol, and 25 μg/mL spectinomycin.

In each conjugation experiment, 2 mL of *Anabaena* cells in the logarithmic phase (OD_730_ = 0.5-0.7) was cultured. Simultaneously, 2 mL of the 481 strain containing the plasmids pCpf1b-sp2, pRL443, and pRL623 (OD600=0.5) was prepared. Both the *Anabaena* and 481 cells were washed 2-3 times using either BG11. Following the washes, a volume of 0.5 mL of the appropriate broth was used to resuspend the *Anabaena* and 481 cells in a sterile cell culture tube. The cell mixture was then incubated at a temperature of 30 °C under light conditions of 30 μE/m2/s for a duration of 4-6 hours. Subsequently, a volume of 0.2 mL of the resuspended cell mixture was transferred onto a BG11 agar plate containing 1.2% agar. A sterile nitrocellulose membrane (NC) was placed over the plate for cover.

The plates were incubated for 1-2 days and then transferred to new BG11 agar plates containing either 5 μg/mL spectinomycin. After incubating the conjugates for 1-2 weeks, they exhibited single colonies. A single colony was selected and streaked onto a new antibiotic BG11 agar plate. The selected colony was then inoculated into BG11 broth supplemented with either 5 μg/mL spectinomycin. After 4-5 days, the *Anabeana* cells carrying the editing plasmid were confirmed by colony PCR and Sanger sequencing.

### Generation of mutant cyanophages

A volume of 300 μL of *Anabeana* cells with an OD_730_ = 0.5 was co-cultured with 100 μL of cyanophage A-1(L) or A-4(L) in 2 mL of BG11 broth at a temperature of 30 °C for a duration of 2-3 days. After co-cultivation, the lysate was centrifuged at 7000 rpm for 3 minutes, and the resulting supernatant was stored at 4 °C. This supernatant served as the wild-type (WT) A-1(L) or A-4(L) sample for use in this study. To generate mutant cyanophages, 100 μL of WT A-1(L)/A-4(L) cyanophage were propagated on 300 μL of *Anabeana* cells carrying editing plasmids (OD_730_ = 0.5) that were grown in BG11 broth supplemented with antibiotics. Subsequently, 500 μL of the supernatant containing the mutant cyanophages was transferred to a fresh culture of *Anabeana* cells containing editing plasmids. This process was repeated for several rounds. The remaining supernatant from each transfer was stored at 4°C for further analysis.

The mutant cyanophages were purified three times. For each purification, dilutions of the cyanophage lysate were mixed with 600 μL of WT Anabeana cells. Then, 10 mL of 0.7% BG11 agar was added and the mixture was plated onto plates to allow for the formation of single plaques. The identity of the single plaques of mutant cyanophages was confirmed through colony PCR and Sanger sequencing. All primer oligonucleotides used in this study were synthesized by Sangon Biotech and are listed in Table S1.

### Characterization of mutant cyanophages

To determine whether the mutant cyanophages differed from the WT cyanophages, we evaluated their one-step growth curve and their ability to inhibit cyanobacterial growth.

To determine the mutant *cyanophages’* ability to infect the host strain, we evaluated their inhibition of cyanobacterial growth using a multiplicity of infection (MOI) of 0.01. Fresh *Anabeana* cells were diluted to an OD_730_ of 0.2, and 400 μL of cyanophage was added to the cell culture tube with 4 mL of Anabeana cells. BG11 liquid served as a negative control, while *Anabeana* cells without cyanophage served as the positive control. The mixture was cultured at 30□ and 120 rpm. We took 200 μL of the mixture every two hours for A-4(L) and six hours for A-1(L) to monitor the OD_730_ using a 96-well tissue culture plate and a microplate reader. Each group was replicated three times.

## Acknowledgments

We would like to extend our gratitude to Dr. Chengcai Zhang from the Institute of Hydrobiology, Chinese Academy of Sciences, for generously providing *Anabeana* sp. PCC.7120, as well as the plasmids pCpf1b-sp, and the cyanophages A-1(L) and A-4(L).

## Supplementary Information

**Supplementary Fig. 1** Maps of plasmids pCpf1b-sp2 used in this study

**Supplementary Fig. 2** Genome maps of the cyanophage A-4 (L)

**Supplementary Fig. 3** Genome maps of the cyanophage A-4 (L)

**Supplementary Table 1**. Gene annotation of the cyanophage A-1 (L)

**Supplementary Table 2**. Gene annotation of the cyanophage A-4 (L)

**Supplementary Table 3**. Part primers used in this study

