## Supplementary_information for "CRISPR/Cas-based genome editing for cyanophage of *Anabeana sp*"

**Figures**

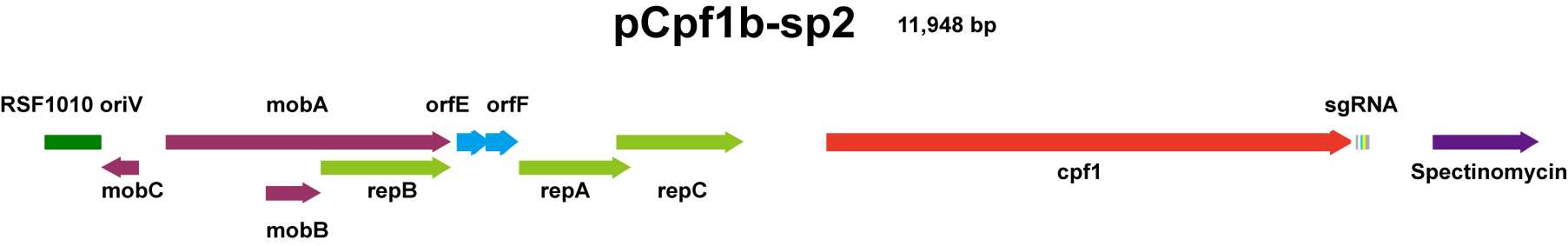

**Supplementary Fig. 1** Maps of plasmids pCpf1b-sp2 used in this study.

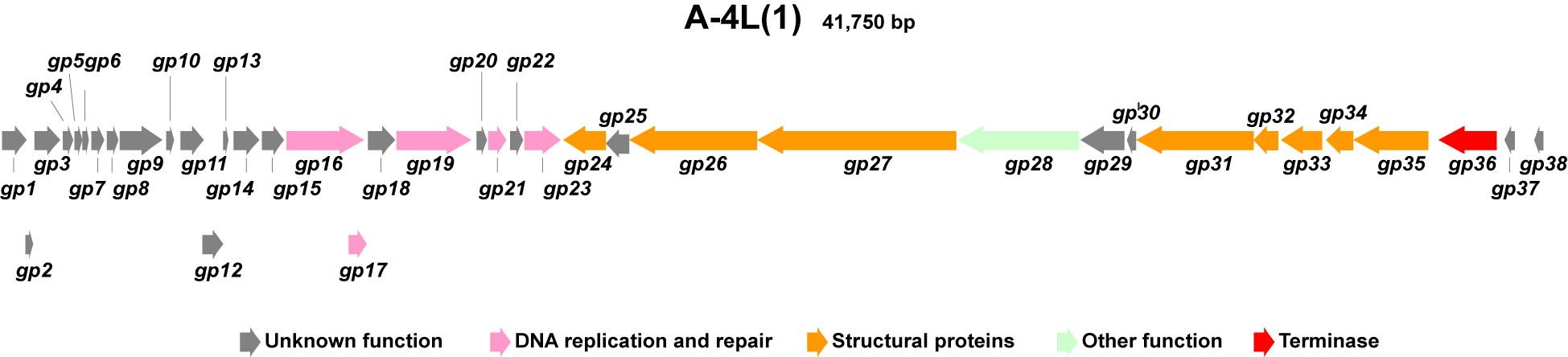

**Supplementary Fig. 2** Genome maps of the cyanophage A-4 (L). Arrows represent the genes and are colored according to the genes’ predicted functions.

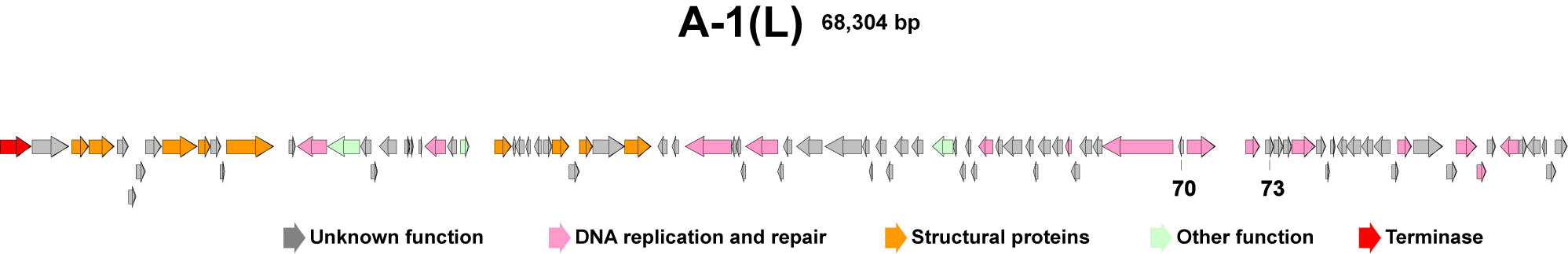

**Supplementary Fig. 3** Genome maps of the cyanophage A-4 (L). Arrows represent the genes and are colored according to the genes’ predicted functions.

**Tables**

**Supplementary Table 1.** Gene annotation of the cyanophage A-1 (L)

| gene_id | start | end | strand | function |
| --- | --- | --- | --- | --- |
| *gp_01* | 432 | 1082 | + | hypothetical protein |
| *gp_02* | 1046 | 1249 | + | hypothetical protein |
| *gp_03* | 1270 | 1953 | + | hypothetical protein |
| *gp_04* | 2028 | 2288 | + | hypothetical protein |
| *gp_05* | 2325 | 2510 | + | hypothetical protein |
| *gp_06* | 2516 | 2698 | + | hypothetical protein |
| *gp_07* | 2756 | 3082 | + | hypothetical protein |
| *gp_08* | 3155 | 3460 | + | hypothetical protein |
| *gp_09* | 3513 | 4610 | + | hypothetical protein |
| *gp_10* | 4724 | 4921 | + | hypothetical protein |
| *gp_11* | 5087 | 5674 | + | hypothetical protein |
| *gp_12* | 5664 | 6185 | + | hypothetical protein |
| *gp_13* | 6181 | 6318 | + | hypothetical protein |
| *gp_14* | 6462 | 7151 | + | hypothetical protein |
| *gp_15* | 7209 | 7784 | + | hypothetical protein |
| *gp_16* | 7830 | 9866 | + | DNA primase/helicase |
| *gp_17* | 9469 | 9936 | + | endonuclease |
| *gp_18* | 9977 | 10660 | + | hypothetical protein |
| *gp_19* | 10692 | 12647 | + | DNA polymerase |
| *gp_20* | 12780 | 13073 | + | hypothetical protein |
| *gp_21* | 13114 | 13554 | + | Erf-like ssDNA annealing protein |
| *gp_22* | 13687 | 13989 | + | hypothetical protein |
| *gp_23* | 14053 | 14979 | + | Cas4-domain exonuclease |
| *gp_24* | 15044 | 16159 | - | tail fiber protein |
| *gp_25* | 16164 | 16769 | - | hypothetical protein |
| *gp_26* | 16769 | 20110 | - | tail fiber protein |
| *gp_27* | 20119 | 25269 | - | tail protein |
| *gp_28* | 25324 | 28497 | - | peptidase |
| *gp_29* | 28513 | 29652 | - | hypothetical protein |
| *gp_30* | 29749 | 29973 | - | hypothetical protein |
| *gp_31* | 29976 | 33020 | - | tail protein |
| *gp_32* | 33023 | 33673 | - | tail protein |
| *gp_33* | 33756 | 34817 | - | major head protein |
| *gp_34* | 34909 | 35622 | - | head scaffolding protein |
| *gp_35* | 35612 | 37570 | - | head-tail adaptor |
| *gp_36* | 37859 | 39346 | - | terminase |
| *gp_37* | 39551 | 39817 | - | hypothetical protein |
| *gp_38* | 40333 | 40587 | - | hypothetical protein |

**Supplementary Table 2.** Gene annotation of the cyanophage A-4 (L)

| gene_id | start | stop | strand | function |
| --- | --- | --- | --- | --- |
| *gp_01* | 1388 | 2992 | + | hypothetical protein |
| *gp_02* | 3146 | 3877 | + | putative outer membrane protein |
| *gp_03* | 3883 | 4980 | + | putative major capsid protein |
| *gp_04* | 5119 | 5607 | + | hypothetical protein |
| *gp_05* | 5604 | 5930 | + | hypothetical protein |
| *gp_06* | 5930 | 6328 | + | hypothetical protein |
| *gp_07* | 6325 | 7047 | + | hypothetical protein |
| *gp_08* | 7079 | 8599 | + | tail sheath protein |
| *gp_09* | 8656 | 9159 | + | tail tube protein |
| *gp_10* | 9171 | 9611 | + | hypothetical protein |
| *gp_11* | 9601 | 9813 | + | hypothetical protein |
| *gp_12* | 9883 | 11,952 | + | tail protein |
| *gp_13* | 12,605 | 12,892 | + | hypothetical protein |
| *gp_14* | 12,978 | 14,255 | - | peptidase |
| *gp_15* | 14,261 | 15,676 | - | lysozyme |
| *gp_16* | 15,700 | 16,167 | - | hypothetical protein |
| *gp_17* | 16,166 | 16,447 | + | hypothetical protein |
| *gp_18* | 16,547 | 17,284 | - | hypothetical protein |
| *gp_19* | 17,651 | 17,848 | + | hypothetical protein |
| *gp_20* | 17,851 | 18,030 | + | hypothetical protein |
| *gp_21* | 18,260 | 18,400 | + | hypothetical protein |
| *gp_22* | 18,533 | 19,423 | - | exonuclease RNAse T and DNA polymerase |
| *gp_23* | 19,497 | 19,934 | - | hypothetical protein |
| *gp_24* | 20,061 | 20,465 | + | LurX regulator protein |
| *gp_25* | 21,549 | 22,307 | + | baseplate assembly protein |
| *gp_26* | 22,363 | 22,470 | - | hypothetical protein |
| *gp_27* | 22,482 | 22,856 | - | hypothetical protein |
| *gp_28* | 22,923 | 23,141 | - | hypothetical protein |
| *gp_29* | 23,289 | 23,618 | - | hypothetical protein |
| *gp_30* | 23,714 | 24,058 | + | hypothetical protein |
| *gp_31* | 24,081 | 24,818 | + | baseplate tail protein |
| *gp_32* | 24,809 | 25,252 | + | hypothetical protein |
| *gp_33* | 25,252 | 25,827 | + | tail collar protein |
| *gp_34* | 25,849 | 27,237 | + | hypothetical protein |
| *gp_35* | 27,253 | 28,392 | + | tail collar protein |
| *gp_36* | 28,664 | 29,092 | - | hypothetical protein |
| *gp_37* | 29,286 | 29,594 | - | hypothetical protein |
| *gp_38* | 29,868 | 31,874 | - | DNA polymerase |
| *gp_39* | 31,901 | 32,092 | - | hypothetical protein |
| *gp_40* | 32,095 | 32,307 | - | hypothetical protein |
| *gp_41* | 32,307 | 32,516 | - | hypothetical protein |
| *gp_42* | 32,500 | 33,915 | - | helicase |
| *gp_43* | 33,912 | 34,142 | - | hypothetical protein |

**Supplementary Table 2. (Continuous)**

| gene_id | start | stop | strand | function |
| --- | --- | --- | --- | --- |
| *gp_44* | 34,139 | 34,504 | - | hypothetical protein |
| *gp_45* | 34,733 | 35,821 | - | hypothetical protein |
| *gp_46* | 35,972 | 37,564 | - | hypothetical protein |
| *gp_47* | 37,604 | 37,879 | - | hypothetical protein |
| *gp_48* | 37,879 | 38,073 | - | hypothetical protein |
| *gp_49* | 38,189 | 38,641 | - | hypothetical protein |
| *gp_50* | 38,631 | 38,915 | - | hypothetical protein |
| *gp_51* | 39,011 | 39,595 | - | hypothetical protein |
| *gp_52* | 39,730 | 40,245 | - | hypothetical protein |
| *gp_53* | 40,664 | 41,554 | - | putative antirepressor |
| *gp_54* | 41,568 | 41,711 | - | hypothetical protein |
| *gp_55* | 41,854 | 42,090 | - | hypothetical protein |
| *gp_56* | 42,087 | 42,341 | - | hypothetical protein |
| *gp_57* | 42,338 | 42,580 | - | hypothetical protein |
| *gp_58* | 42,687 | 43,304 | - | DNA N-6 adenine methyltransferase |
| *gp_59* | 43,398 | 43,688 | - | hypothetical protein |
| *gp_60* | 43,737 | 44,573 | - | hypothetical protein |
| *gp_61* | 44,736 | 45,059 | - | hypothetical protein |
| *gp_62* | 45,046 | 45,264 | - | hypothetical protein |
| *gp_63* | 45,261 | 45,797 | - | hypothetical protein |
| *gp_64* | 45,897 | 46,331 | - | hypothetical protein |
| *gp_65* | 46,462 | 46,704 | - | ASCH domain protein |
| *gp_66* | 46,706 | 47,056 | - | hypothetical protein |
| *gp_67* | 47,053 | 47,616 | - | hypothetical protein |
| *gp_68* | 47,650 | 48,054 | - | hypothetical protein |
| *gp_69* | 48,076 | 51,141 | - | DNA primase |
| *gp_70* | 51,375 | 51,617 | - | hypothetical protein |
| *gp_71* | 51,775 | 52,983 | + | transposase |
| *gp_72* | 54,304 | 54,942 | + | thymidylate kinase |
| *gp_73* | 55,179 | 55,553 | + | hypothetical protein |
| *gp_74* | 55,550 | 55,966 | + | hypothetical protein |
| *gp_75* | 55,976 | 56,335 | + | hypothetical protein |
| *gp_76* | 56,332 | 57,375 | + | DNA-cytosine methyltransferase |
| *gp_77* | 57,407 | 57,820 | + | hypothetical protein |
| *gp_78* | 57,820 | 57,987 | + | hypothetical protein |
| *gp_79* | 58,019 | 58,234 | + | hypothetical protein |
| *gp_80* | 58,298 | 58,708 | - | hypothetical protein |
| *gp_81* | 58,722 | 59,327 | - | hypothetical protein |
| *gp_82* | 59,379 | 59,885 | - | hypothetical protein |
| *gp_83* | 59,898 | 60,623 | - | hypothetical protein |
| *gp_84* | 60,685 | 60,987 | + | hypothetical protein |
| *gp_85* | 60,950 | 61,549 | + | dCTP deaminase |

**Supplementary Table 2. (Continuous)**

| gene_id | start | stop | strand | function |
| --- | --- | --- | --- | --- |
| *gp_86* | 61,635 | 62,933 | + | hypothetical protein |
| *gp_87* | 63,072 | 63,518 | + | hypothetical protein |
| *gp_88* | 63,508 | 64,395 | + | DNA methylase |
| *gp_89* | 64,385 | 64,816 | + | RusA |
| *gp_90* | 64,866 | 65,210 | + | hypothetical protein |
| *gp_91* | 65,435 | 66,214 | - | thymidylate synthase |
| *gp_92* | 66,258 | 66,572 | + | hypothetical protein |
| *gp_93* | 66,575 | 67,171 | - | hypothetical protein |
| *gp_94* | 67,221 | 67,421 | - | hypothetical protein |
| *gp_95* | 67,455 | 67,838 | + | hypothetical protein |
| *gp_96* | 67,816 | 68,304 | + | hypothetical protein |
| *gp_97* | 1 | 1358 | + | terminase large subunit; TerL |

**Supplementary Table 3.** Part primers used in this study

| primers | primer sequence |
| --- | --- |
| a1.F | GAGACCACTAGTGGTCTCAatctacaacagtagaaattatttaaagttcttagacccg |
| s2.R | tttgcttccagtcgtttggtatggcttcattcagct |
| a2.F | accaaacgactggaagcaaagccaggaaagcgg |
| s1.R | tagcgatttatgaaggtcatttttttgtctagct |
| backbone.F | cttgctcaatctgaaagcgaccaggtgctcg |
| backbone.R | gtcgctttcagattgagcaagctttatgcttgtaaaccg |
| laz-a.F | caacgttgttgccattgcgga |
| laz-s.R | agaattcttgacagctagctcagtcc |
| scb-a.F | ttatgcagtgctgccataaccatgag |
| scb-s.R | aaaaattgcctactgagcgctgc |
| A4-2.test.F | cctgttgctagtgatagcccatc |
| A4-2.test.R | cttgttgaatatccagagcgttagc |
| A4-3.test.R | gcgaatggtactgtgttggttagc |
| A4-5.test.F | gtattgtggctggtgtaccactg |
| A4-5.test.R | ggttagtagggtatgtttgataggtg |
| A4-8.test.F | gaagatgttaggctcactacttctcatgg |
| A4-8.test.R | gggttaatgttcgtctcatcgtgg |
| A4-18.test.F | gaagcgaagggcttattcagtgc |
| A4-18.test.R | actcccatgtctctgtccaactg |
| A1-70.test.F | ttgtttttggtttccggtgttg |
| A1-70.test.R | agttttaatcgcctcatccagc |
| A1-73.test.F | cccgattgcagtttttaggaatg |
| A1-73.test.R | actgcgaagtaccaaaaattcaaattg |
| gRNA.test.F | atgaagagtattttgagttcgtgcag |
| gRNA.test.R | gcaaggtttcggtcttctagagtc |
